# *Enviromics* in breeding: applications and perspectives on envirotypic-assisted selection

**DOI:** 10.1101/726513

**Authors:** Rafael T. Resende, Hans-Peter Piepho, Orzenil B. Silva-Junior, Fabyano F. e Silva, Marcos Deon V. de Resende, Dario Grattapaglia

## Abstract

Genotype by Environment interaction (G × E) studies have focused mainly on estimating genetic parameters over a limited number of experimental trials. However, recent Geographic Information System (GIS) techniques have opened new frontiers for understanding and dealing with G × E. These advances allow increasing selection accuracy across all sites of interest, including those where experimental trials have not yet been deployed. Here, we introduce the term Enviromics under an envirotypic-assisted breeding framework and propose the GIS-GE method, i.e. a geospatial tool to maximize genetic gains by predicting the phenotypic performance of unobserved genotypes using “enviromic markers”. In summary, a particular site represents a set of envirotypes, each one representing a set of environmental factors that interact with the genetic background of genotypes, thus resulting in informative re-rankings to make decisions over different environments. Based on a simulated case study, we show that GIS-GE allows accurate (*i*) matching of genotypes to their most appropriate sites; (*ii*) definition of breeding areas that have high genetic correlation to ensure selection gains across environments; and (*iii*) indication of the best sites to carry out experiments for further analysis based on environments that maximize heritability. Envirotyping techniques provide a new class of markers for genetic studies, which are inexpensive, increasingly available and transferable across species. We envision a promising future for the integration of the Enviromics approach into breeding when coupled with next-generation genotyping/phenotyping and powerful statistical modeling of genetic diversity. Environmental scenarios can also be improved using information from strategic plans for biodiversity and genetic resources management, especially in the current perspective of dynamic climate change.

**Key message:** We propose the application of *Enviromics* to breeding practice, by which the similarity among sites assessed on an “omics” scale of environmental attributes drives the prediction of unobserved genotypes.

## Introduction

One of the greatest challenges of modern agriculture is dealing with the limited prospects of significantly expanding farmed land. Tailoring highly adapted genetic material to the available environments becomes a key element to increase agricultural yields without the conversion of additional land and losses due to adverse environmental impact (Garnett et al. 2013). The differential response of genotypes across variable environments, known as Genotype by Environment (G × E) interaction, represents one of the major challenges faced by essentially all breeding programs. Traditional G × E studies are based on the restricted evaluation of a few trials designed to find estimates of parameters that capture the genotypic interaction of the tested genetic materials with the environments (Elias et al. 2016).

Phenotypic variation results from genetic and environmental influences, which have to be taken into account in the evaluation of phenotypic plasticity (Nicotra et al. 2010). To maximize genetic gains, it is thus necessary to collect data from all measurable environmental factors affecting the performance of genotypes in any particular site (Des Marais et al., 2013). Patterns of relationships between environmental variables and the expression of genotypes can be exploited through modern concepts of phenomics and genomics, which rely on high-throughput, large scale data collection and evaluation techniques (Houle et al. 2010; van Eeuwijk et al. 2018). Even here, the changes in the physical environment have not yet been well exploited in the genetic analyses that seek correlations between environmental factors and yield or performance variables.

Much of the actual research in this area at the interface of genetics and breeding focuses on building options for selection against the genotype’s environmental sensitivity (Rauw and Gomez-Raya 2015). Usually, these approaches rely on the separation of different sources of variability contributing to the genotype’s performance by looking at the changes of the values for a particular trait in a new set of conditions in different environments. The quantity of change, in terms of the variance of the normal population from which the genotype is sampled, is used to ascribe to the different causes of variability in the genotype’s response due to its genetics, to the environment and the fraction which arises from the interaction of a particular hereditary makeup with a particular kind of environment. In advanced genetic analyses, the total patterns of expression of a trait among the environments by a genotype or by various genotypes along descriptors of the environment, are analyzed using reaction norm models (De Jong 1995), which rely on the definition of an environmental variable. In breeding practice, an environmental variable for a reaction norm model of selection is usually calculated as the mean phenotypic performance of a trait in a restricted range of environments (Finlay and Wilkinson 1963; Eberhart and Russell 1966) and genetic covariance structures are proposed to re-rank breeding values including Genotype by Environment interaction (Calus et al. 2004). The effect of describing an environmental variable when building genetic covariance structures for reaction norm models of selection has been currently investigated in animal breeding (for a review see Rauw and Gomez-Raya, 2015).

The possibility of using experimental data from a particular environment to anchor the prediction of performance and subsequent recommendation of potentially successful genotypes in other untested sites has been a theme of great interest in plant breeding, explored by a number of authors by addressing environmental similarity based on multiple environmental attributes. To be able to predict the performance of individuals in an untested site, environmental covariables need to be used (Piepho et al. 1998). The combination of environmental covariables with Geographic Information Systems (GIS), was also proposed early on by Annicchiarico et al. (2006). The use of extensive environmental information in reaction norm models was recommended by Jarquín et al. (2014). These last authors used 68 environmental covariates to evaluate grain yield of commercial wheat lines, allowing the prediction of unobserved lines in untested environments based on exploiting high-dimensional environmental and genomic data. Several other authors have implemented and expanded this idea, such as Pérez-Rodríguez et al. (2015) who used 76 environmental covariables and pedigree information in nine cotton trials. Additionally, to obtain greater predictive accuracy through the exploration of environmental co-variation, the concept of on-farm trials has been used, using large volumes of data to tap into hundreds or even thousands of environments (Hernández et al. 2018). The inclusion of environmental information in the genetic models has resulted in significant gains in accuracy, improving the predictive ability by 20% and, in some cases, by up to 34% (Jarquín et al. 2014; Acosta-Pech et al. 2017). Prediction using environmental data also opens possibilities to expand recommendations of genetic materials across countries that lack experiments for particular crops of interest, thus saving resources, avoiding sanitary barriers and reducing phenotyping costs (Pérez-Rodríguez et al. 2017; Sukumaran et al. 2017).

Model types such as mixed-effects, linear-bilinear, crop growth and Bayesian ones have shown their consolidated strength in analyzing data from multiple environments (van Eeuwijk et al. 2016). Notwithstanding the impact that such approaches have had in understanding and exploiting the G × E interaction for prediction of yet-to-be-observed phenotypes, current models do provide room for expanding their use to features not yet explored. Understanding the sources of environmental variation has increasingly become a key element for the assessment and recommendation of genotypes under probabilistic scenarios of global climatic changes and rapid landscape modification by human actions (Raza et al. 2019). Such models can be used along the stages of a breeding cycle (Annicchiarico and Iannucci 2008) to identify loci related to phenotypic trait expression (Gauch et al. 2011) and to determine sets of environmental factors underlying phenotypic plasticity (Piepho 2000; Nicotra et al. 2010). Among the reported models, genomic selection (GS) and genome-wide association studies (GWAS) can also be developed under environmental gradient models (Acosta-Pech et al. 2017; Grattapaglia et al. 2018).

In this study, we explore the concept of *Enviromics* in the context of plant breeding by presenting and applying a method based on Geographic Information Systems coupled with genetics (GIS-GE) to a case study based on advanced environmental interpolation techniques. We show that the main advantages brought about by this method are an improved matching of genotypes to their most appropriate sites, the improved zoning of breeding areas with high genetic correlation, and the indication of the best sites to carry out experiments for further analysis based on regions that maximize the heritability.

### Background integrating Enviromics with breeding

To explore more thoughtfully the influences of the various environmental factors on selective breeding, we borrow the term “Enviromics” – all the environmental conditions that affect human health in the scope of precision medicine (Gad 2008; Riggs et al. 2018) – and extend it to the environment-dependent part of reaction norm models for selection in the perspective of exploiting patterns of G × E in local environments, mainly in plant, but promptly extendable to animal breeding. This work is inspired by the exciting developments in the field of population genomics and epidemiology in which a new type of analysis of phenotypic and genotypic data and environmental variables, termed “phenome-wide association studies”, have been used to identify effects of environmental variables on clinical characteristics using data from large and diverse populations such as the PAGE consortium (Matise et al. 2011). Along these same lines, a recent study involving an extensive analysis of the local environments described by 204 geoclimatic variables of *Arabidopsis* accessions and 131 phenotypes, revealed candidate adaptive genetic variation such as cold tolerance associated with high-dimensional environmental variables (Ferrero-Serrano and Assmann 2019).

In our conceptualization of Enviromics in breeding, a particular land area is a geoprocessing environment corresponding to a grid of pixels, just like those of a digital photograph, and for any single environmental variable a value can be assigned to every pixel (see the top of Figure 1). The value distribution of a particular environmental variable in this collection of pixels constitutes the envirotypes that have developed in the land area. In plants, where propagation systems permit obtaining large numbers of identical copies of the same genotype, a particular individual genotype can be represented by an inbred line, hybrid or clonal variety that is tested in multiple sites so that the notion of a reaction norm of the genotypes spread across a wide environmental range is quite direct. In animal breeding, the reaction norm of an individual genotype (often the sire) needs to be approximated by the performances of its offspring across a range of environments (Rauw and Gomez-Raya 2015). Since phenotypic data from an individual genotype in existing field trials is available in some pixels, one can relate the distribution of a particular phenotypic value with the envirotypes using statistical modeling (Hyman et al. 2013; van Eeuwijk et al. 2018). The environmental information that can be obtained by *Envirotyping* (Xu 2016) can be termed “enviromic markers”. In turn, the association of all these markers composes the *envirome*. Envirotyping is a third “typing” technology, alongside genotyping and phenotyping. Enviromics is a corresponding third “omics” technology, complementing genomics and phenomics technologies. The possible facets that an enviromic marker presents within site/pixel are the envirotypes. The marker polymorphism is the envirotypic variation. Reaction norm models are then built for the whole evaluated area and used to predict the performance of any tested genotype for the trait under evaluation in any pixel (tested and non-tested) in the geoprocessing environment.

**Figure 1.**
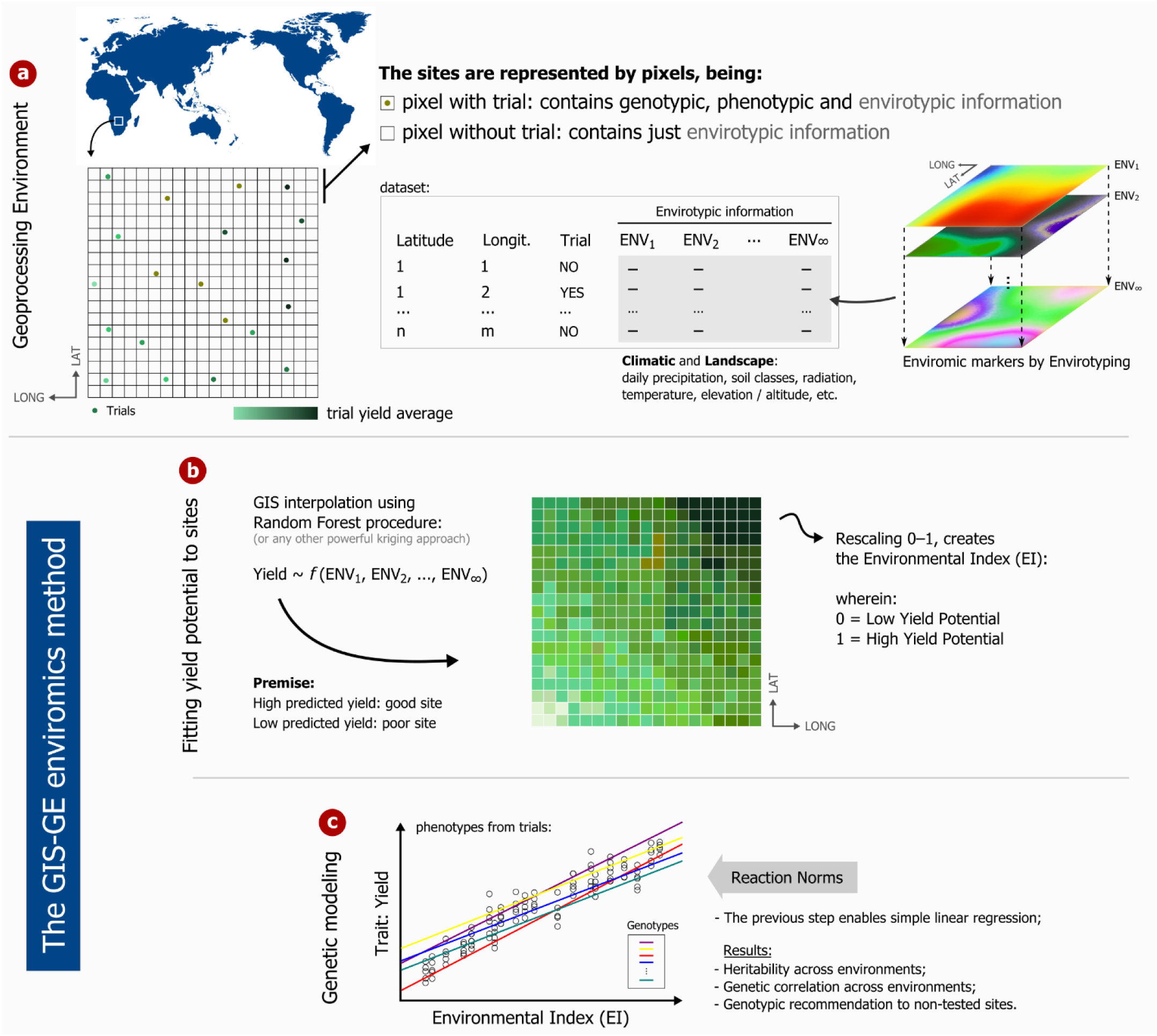
**(a)** Example of an Enviromics data set and **(b and c)** a hypothetical GIS-GE method application containing just 20 trials and 5 genotypes.

Integrating breeding and environmental data relies on the increasing worldwide availability of geoprocessing technologies, such as GIS in the scope of Precision Agriculture (PA) (Lindblom et al. 2017). The collection and processing of spatio-temporal data on climate, water, soil and yields variables is rapidly increasing due to societal need for food security and technological advances. The Arabidopsis CLIMtools repository (http://www.personal.psu.edu/sma3/CLIMtools.html) is, to our knowledge, the first repository of data and tools tailored for the exploitation of environmental variation associated to any gene or variant of interest in plants. In crop plants, the international DivSeek initiative is one remarkable example (Nature Genetics Editorial 2015) and we can envisage in the near future a rapid expansion of similar efforts toward the organization and availability of very large records of genetics and genomics data linked to physical resources to germplasm curators, breeders and researchers alike.

The main challenge in the implementation of Enviromics approaches is that only a few of the pixels representing a land area contain breeding trials having phenotypic and genotypic information. However, all pixels have environmental information since affordable meteorological stations have been increasingly installed everywhere, such that environmental data can be interpolated by kriging across any desired area (Oliver and Webster 2015). Because geospatial information is now easily accessible, data-driven approaches supported by GIS can be exploited for breeding practice. The challenges of data analysis in agricultural research are changing as the features of available data improve (Wolfert et al. 2017), especially through the use of modern geotechnologies (Xu 2016). Remote-sensing technologies can also provide data that can be used with specific spectral bands for the aimed purposes (Kasampalis et al. 2018). Precipitation, temperature, terrain altitude (in the form of digital elevation models), some cultural treatment, solar radiation, soil properties and water deficit information are some of the variables that relate well to most phenotypic traits evaluated in agricultural production (Xu 2016; Chang 2017). Indeed, the partnership between GIS and breeding has been gradually showing solid foundations (Hyman et al. 2013; Haghighattalab et al. 2017; Marcatti et al. 2017). The spatial positioning of trials or even experimental plots linked to the measurement of multiple environmental information by envirotyping, either at macro or micro scales, suggests that the adherence of enviromic techniques is especially promising in association with breeding cycles and genotypic recommendations.

## Methods

### Environmental and genetic data simulation

We simulated a didactical dataset with 50 experimental trials comprising 100 plant genotypes named G001, G002, up to G100, using R software (Team R 2015). In the simulation, two sequential stages were adopted. The first consisted of the land area composition into a geoprocessing environment and formulation of the envirotypic data, independent of genotypic and phenotypic information. The second one consisted of generating trials and genotypes, while the plant phenotypes were built in such a way as to have four variant sources: (i) genotypic values normally distributed with known mean and variance, (ii) a genotypic relationship with the envirotypic information of the previous simulation stage, (iii) a particular trial effect, and (iv) an overall random error. Figure 1-a shows a schematic example containing some trials, and the arrangement of the envirotypic data of each enviromic marker.

The 50 trials were randomly allocated to a square area covering 10,000 pixels. The mimicked response trait was agricultural yield, which could correspond, for example, to the yield of an annual grain crop, timber volume of a planted forest tree, forage biomass or fruit yield of a fruit crop. We simulated data for external environmental variables that could potentially affect yield in the area. One hundred overlapping *rasters* containing envirotypic information were simulated, each *raster* corresponding to a grid of all pixels in the area, i.e. an enviromic marker. In our simulation these external environmental variables were merely random numbers, but in practice they would correspond, among others, to mean annual temperature, historical annual precipitation, some cultural treatments, terrain altitude, soil characteristics, radiation and vegetation indices, as described by Xu (2016), when classifying external environmental variables. These enviromic markers simulated contribute to the phenotypic trait expression either positively or negatively. Their influence was assumed to be latent, i.e., the size of the effects of each of them on the phenotypic trait were not registered or saved along the simulations. The simulation process was repeated countless times until the broad-sense heritability was approximately 10% (*h*^2^ ≈ 0.10) aiming at a representation of a typical quantitative trait. To consign this, the model used was as described below in the ‘Genetic modeling’ section by Eq. [1]. The phenotypic mean was 46.10 units with standard deviation equal to 12.90 and maximum and minimum values equal to 5.50 and 117.10, respectively. To illustrate this simulation framework, three enviromic markers as well as the spatial distribution of the genotypic trials within the target area are shown in Figure 1a.

### Case Study: GIS-GE method for Enviromics

The geospatial genetics (GIS-GE) method proposed here allows for a joint analysis of the experimental setting accounting for phenotypic, genotypic and envirotypic data. The procedure generates a map with optimal recommendation of genotypes to identify geographic zones with high genetic correlation between environment variables and individual genotypes, i.e., locations in which rank changes of genotypes are small. Additionally the method identifies locations that maximize the accuracy of genetic models, allowing an improved capture of trait heritability due to G × E which, in turn, improves selective accuracy. The GIS-GE methodology advances by introducing a new way of evaluating the phenotypic trait by converting the land area into a set of pixels in a geoprocessing framework, making full use of the envirotyping implementation for breeding purposes. The analytical procedures used in GIS-GE consist of two steps. The first one is the development of the enviromic markers relating the trait and environmental variables based on which an Environmental Index (EI) is built. The second one is the genetic modeling, which consists in fitting reaction norm mixed models assuming the EI as an independent variable. This second step also considers the estimation of genetic-environmental parameters such as trait heritability based on the enviromic markers. These procedures are summarized in Figure 1 (parts b and c) and detailed below.

#### Step 1: Enviromental Index (EI)

The phenotypic mean within the 50 experimental trials deployed in the area is calculated and we subsequently generate values of this variable for the entire range of pixels of the raster by using a Random Forest (RF) regression in R software (Liaw and Wiener 2002), a nonparametric multivariate modeling technique that is well suited to capture nonlinear dependencies that uses a common machine learning algorithm based on an enhanced utilization of *regression trees*. According to Koch et al. (2019), several studies from different geoscience fields have shown that RF has overcome most other machine learning techniques available at the time. In our study, five hundred decision trees (default arguments of the *randomForest* R function) were built to establish the relationship between the mean performance of the genotypes within trial for the evaluated yield trait and the 100 enviromic markers. Therefore, the RF adjusted from the phenotypic mean data are used to predict the Environmental Index (EI) across all pixels in the area, using kriging interpolation (see Figure 1-b). We assumed that the higher the predicted phenotypic mean of the site the higher the adaptive fitness, and consequently, the better the site quality. Finally, the EI values were rescaled to 0– 1: 0 being the worst site, and 1 the best one. In addition to composing the genetic model elucidated below, the EI has the important role of imputing the envirotypes for all pixels of the area, which may then be used for further breeding inferences.

Like any other data-based modeling technique, the RF algorithm requires training and validation. The verification of the EI quality for the whole area is carried out by leave-one-out cross-validation (Kohavi 1995), and the model was trained with data of 49 trials and predicted for the 50^th^. The validation procedure is repeated until all the environments have a predicted value, and subsequently the correlation between the observed and predicted values was calculated.

#### Step 2: Genetic modeling

A first model was used considering the trials as experimental blocks, in the shape of:

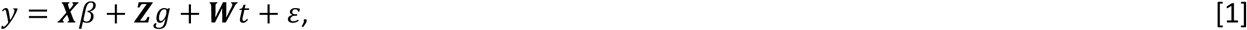

where: **y** is the vector of phenotypic means per genotype and trial ; *β* represents the fixed effects (overall intercept); *g* represents the random effects of genotypes with variance-covariance structure as 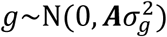; ***A*** is the numerator relationship matrix; *t* represents the random effects of trials with variance-covariance structure as 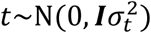; ***X***, ***Z*** and ***W*** are the incidence matrices of *β, g* and *t*, respectively. The residual vector *ε* was assumed as 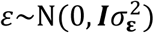. The broad-sense heritability is given as 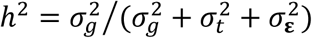.

To represent the association of the evaluated trait with the EI, the following linear model can be adopted (a univariate case) (Resende et al. 2001):

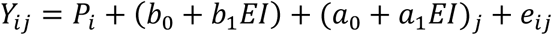

where *Y*_*ij*_ is the measurement of the genotype *j* associated with the fixed effect of the *i*^th^ trial (*i* = 1, 2, …, 50; *j* = 1, 2, ⋯, 100); *P*_*i*_ represents the fixed main effect of the i-th trial; *b*_0_ and *b*_1_ are fixed intercept and slope for EI, the environmental index; *a*_0_ and *a*_1_ are random intercept and slope coefficients of individual *j*, both jointly forming the random effects of additive genetic value; and *e*_*ij*_ is the random error.

Through this model, the genetic value adjusted for a given individual can be seen through two regression groups: fixed regression for all individuals belonging to the same level of fixed effect, which describes the general form for a given individual; random regressions describing the deviations from the fixed regression, generating different slopes for different individuals.

In matrix notation, this model can be written as follows in the Eq. [2]:

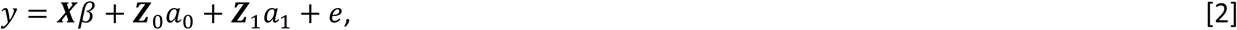

wherein, *y* is the data vector; *β* is the vector of fixed effects; *e* is the vector of random errors; *a*_0_ and *a*_1_ are vectors referring to the coefficients of random regression, which relate the intercepts and EI slopes, respectively. ***X*** is the incidence matrix for fixed effects; ***Z***_0_ is the incidence matrix for *a*_0_, containing 0 and 1’s; ***Z***_1_ is a matrix associating *a*_1_ with *y*, containing zero and EI values. We have the following model averages (math expectations) across trials: *E*(*y*) = ***X****β* and *E*(*y*_*ij*_) = *P*_*ij*_ + (*b*_0_ + *b*_1_ *EI*), wherein (*b*_0_ + *b*_1_ *EI*) refers to the overall mean in EI. The structure of genetic variances is given by

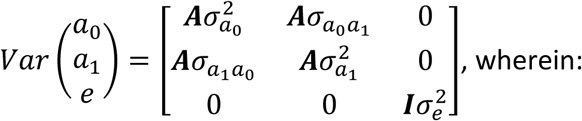

***A*** is the numerator relationship matrix; 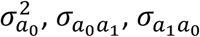 and 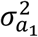 are variances and covariances of and between the random regression coefficients, i.e. they are covariance functions that continuously describe the covariance structure for the trait, in the range of EI covered by the data.

The covariance matrix (***Σ***_*a*_) between random genetic effects for an individual genotype, disregarding kinship relationships, is:

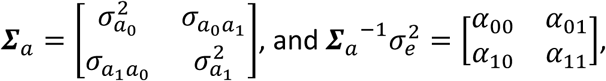

The estimation and prediction of fixed and random effects according to the defined linear model is performed through the mixed model equations:

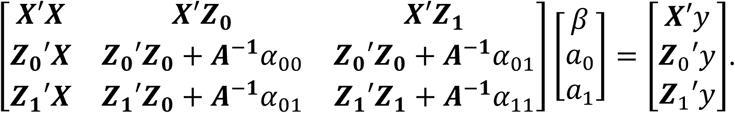

Genetic and residual variances are dependent on EI, that is, they can increase or decrease throughout sites. It can be written as: 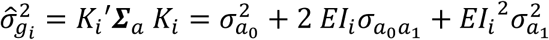 that is the genetic variance in the *i*^th^ EI; *K*_*i*_ a vector of size two, being 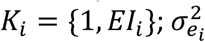 is the residual variance in the *i*^th^ EI, calculated marginally by the model [1], using subsets of 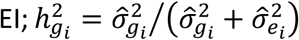, is the heritability for a particular 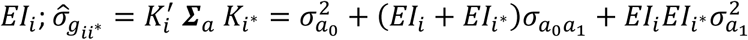 is the genetic covariance between sites *i* and *i*^*^ (addressed by EI information); 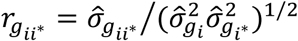 is the genetic correlation between sites *i* and *i*^*^ (addressed by EI information).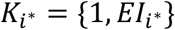.

For the use of mixed model equations and estimation of variances, covariance and genetic correlations, estimates of ***Σ***_*a*_ and 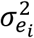 are required. Properties of the genetic parameters estimation for longitudinal data were obtained by restricted maximum likelihood (REML). Based on the EM algorithm we have the following estimators for the components of variance:

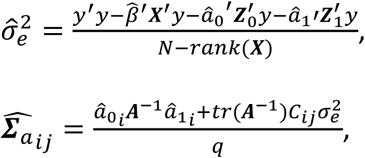

wherein, *N* is the total number of observations. *rank*(***X***): rank of ***X*** or number of columns linearly independent of X; *q*: number of random elements (i.e., number of individuals or genetic values to be predicted). *tr*(): matrix trace operation or sum of matrix diagonal elements. *C*_*ij*_: submatrix that comes from the generalized inverse (*C*) of the matrix of the coefficients of the mixed model equations:

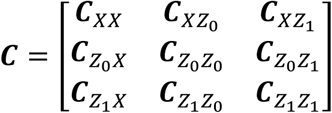

Starting from initial values for ***Σ***_*a*_ and 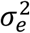, we obtain *β, a* and *λ* from the mixed model equations, which are used in the estimators and 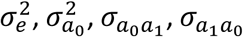 and 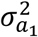 whose estimates are returned to the mixed model equations. This is done successively until convergence. Initial values of 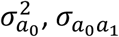 and 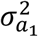 can be obtained from estimates of 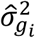 for some EI, by the method of restricted maximum likelihood (REML).

Estimated genetic values (EGV) for genotype *j* can be predicted for various sites (indicated by the EI) through random linear regression: 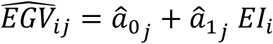. The EGV predictions allow the re-ordering of the candidates to the selection, in the different sites, allowing the re-selection of individuals according to the desired EI.

Based on the genetic correlation matrix among grouped EI (i.e., pixels addressing 0 ≤ EI < 0.1, 0.1 ≤ EI < 0.2, …, 0.9 ≤ EI ≤ 1), we used the UPGMA procedure (Sokal 1958) to define breeding zones, that are locations within which genetic correlations between genotype and EI were optimized. In addition, a recommendation map for potential yield of the best ranked genotypes was also provided as additional visual information derived from GIS-GE.

Finally, in order to evaluate the proposed methodology under unbalanced conditions, the random regression model was tested for the following two situations described below. The success inferences were carried out based on the phenotypic average of the selected genotypes (i.e., the first best genotype ranked for each pixel).

- Randomly reduction of the number of trials (down to a minimum of three). The data from trials that were removed were assigned as validation groups and those remaining as training groups. Thus, more than one draw was made in each reduction stage accounting for approximately a total of 1000 iterations;
- Different levels of genotypic unbalance per trial with the constraint that all genotypes were always present in the analysis, i.e. for each genotype, the number of experiments was reduced (approximately 500 iterations were performed). The data from genotypes that were removed were assigned as validation groups and those remaining as training groups.

## Results

### Development of the Environmental Index

The EI was constructed by extrapolating yield for the whole area, and subsequently rescaling it to 0–1. The mean and EI deviations for all pixels were 0.53 ± 0.27, and for the 50 trials was 0.43 ± 0.29. The distribution of the Environmental Index (EI) for the entire area presented an irregular shape with a higher density around 0.1 and 0.75. Lower densities were observed in the lower and upper tails (Figure 2a). Figure 2b shows the behavior of the 100 genotypes along the range of EI values in terms of the reaction norm for all EIs. The EI distribution for the 50 trials showed a well-distributed range between 0 and 1 (Figure 2c).

**Figure 2.**
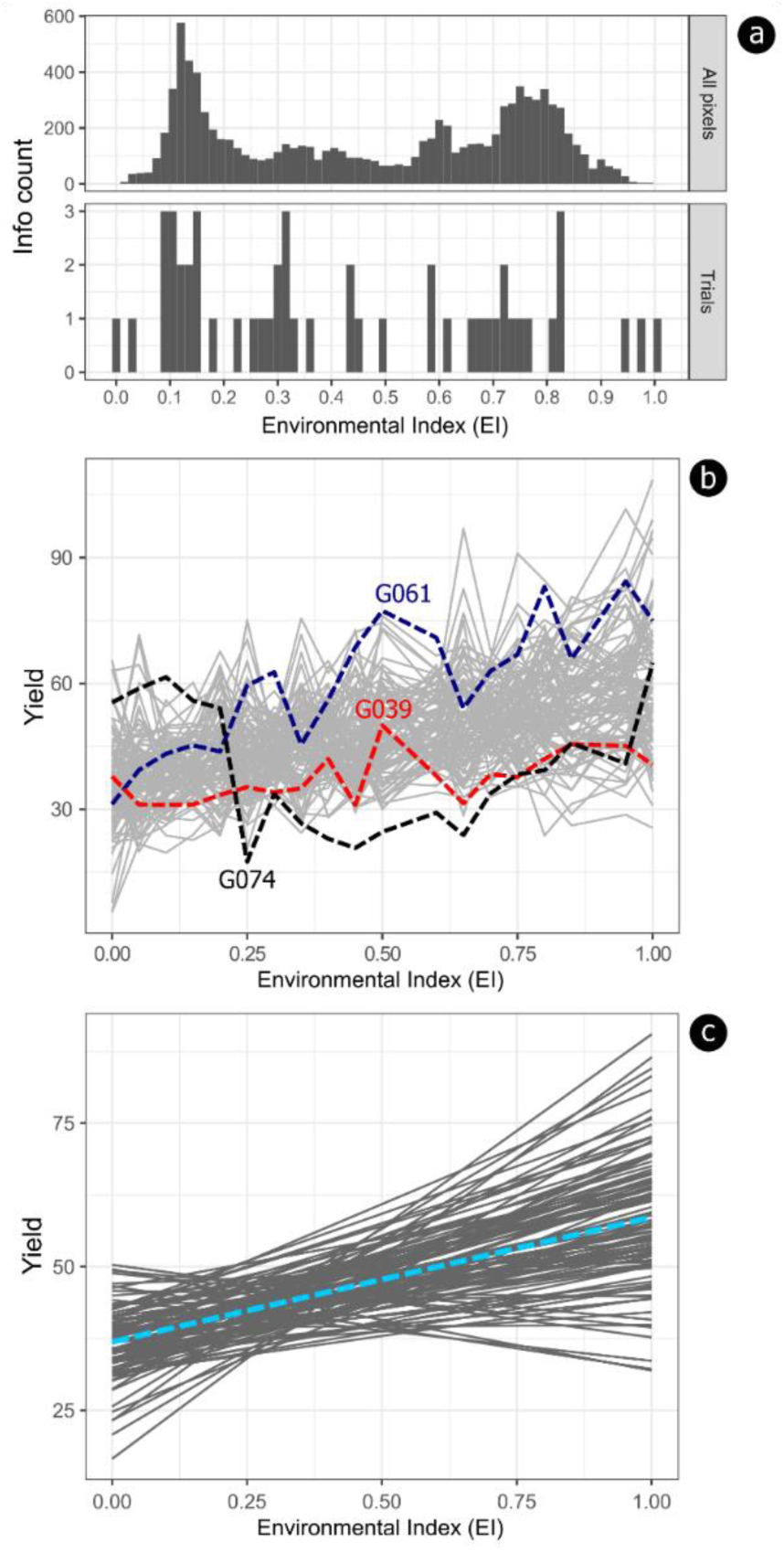
Features of the Environmental Index (EI) obtained in the simulations**. (a)** Comparison between EI of all area pixels and EI of the fifty trials; **(b)** Phenotypic means across environments for the 100 simulated genotypes: genotypes with the highest (G061), lowest (G039) and intermediate (G074) yield are highlighted. **(c)** Reaction norm from the linear random regression model addressing the 100 genotypes. The blue dotted line is the overall fit tendency.

We observed a great overlap between the EI of the whole area and EI subset to the 50 trials (Figure 2a), which is a desired feature for the application of the GIS-GE approach. Moreover, the EI is expected to show a positive correlation with the simulated trait yield. The correlation between the average yield of the 50 trials and the EI was 0.98; whereas it was 0.54 when considering both trials and genotypes. The decreased correlation is explained by the genotypic variability existing within each trial. Following cross-validation, the predictive ability of the RF model was 0.87. These metrics together highlight the suitability of the GIS-GE approach even when there are no yield records from existing trials in the target area. Finally, the general mean and standard deviation of yield considering all 100 genotypes and interpolating to the whole area (10,000 pixels) was equal to 47.23 ± 5.46.

### Genetic modeling

Throughout the pixels of the area, and considering the assumed range of EI (0.00 to 1.00), the broad sense heritabilities varied between 0.41 and 0.47, with the lowest valley value equal to 0.19 in EI = 0.32. We also observed that the two highest heritabilities were seen at the EI extremities (Figure 3a). The average estimated heritability was 0.32, thus larger than the parameter value used in the simulation (without the G × E factor). In terms of the worst (EI=0) and the best (EI=1) environments, the genetic variances were 39.18 and 138.29, respectively. The lowest value was equal to 16.95 at EI = 0.30. The residual variance was 56.75 at (EI = 0) and 154.50 at (EI = 1), so that its lowest value was 56.75 at EI = 0. The model without EI, which considers the trials as experimental blocks presented an overall 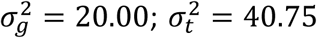, and 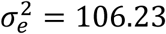, resulting in an overall broad sense heritability equal to 0.12.

**Figure 3.**
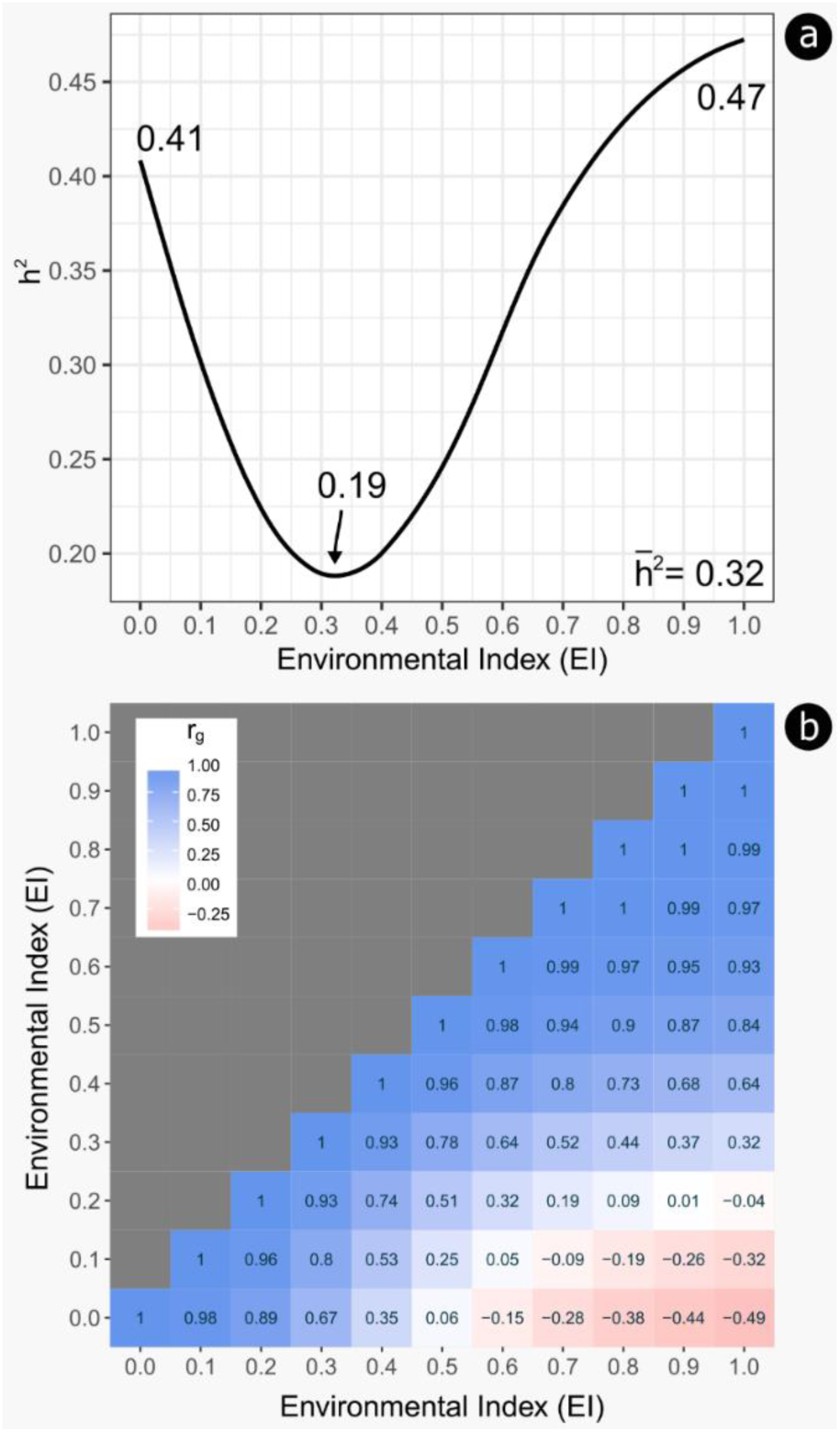
Genetic parameters across the Environmental Indexes. **(a)** Heritability estimate across the entire EI range. **(b)** Heatmap depicting the genetic correlation between different EI values.

Higher genetic correlations (r_g_) were observed between EI=1.00 and EI=0.50, with rg values between 0.99 and 0.84, suggesting low re-ranking between genotypes in these locations (Figure 3b). The lowest genetic correlation laid between the opposites EI=0.00 and EI=1.00, equal to -0.49. The UPGMA procedure for grouping the locations with high genetic correlation between EI and genotypes resulted in the definition of three breeding zones: red (EI between 0.00 and 0.32); khaki (EI between 0.33 and 0.44) and blue (EI between 0.45 and 1.00) (Figure 4a). Among the set of fifty trials, twenty-four belonged to the red zone, four experiments were within the khaki zone, and the blue zone encompassed twenty-one trials. Within the red zone, the genetic correlation measured by rg was, on average, equal to 0.92; within the khaki zone its average value was equal to 0.98; and it was 0.96 within the blue zone. The genetic correlations between the red and khaki zones, red and blue and khaki and blue were respectively 0.39, -0.05 and 0.70, indicating a reasonable genetic re-ranking shared between the khaki and the blue breeding zones. The red zone is the one with the lowest yield potential, with an average environmental index of 0.16, the khaki zone an intermediate potential (on average, EI= 0.38), and the blue zone the highest potential (on average, EI=0.72).

**Figure 4.**
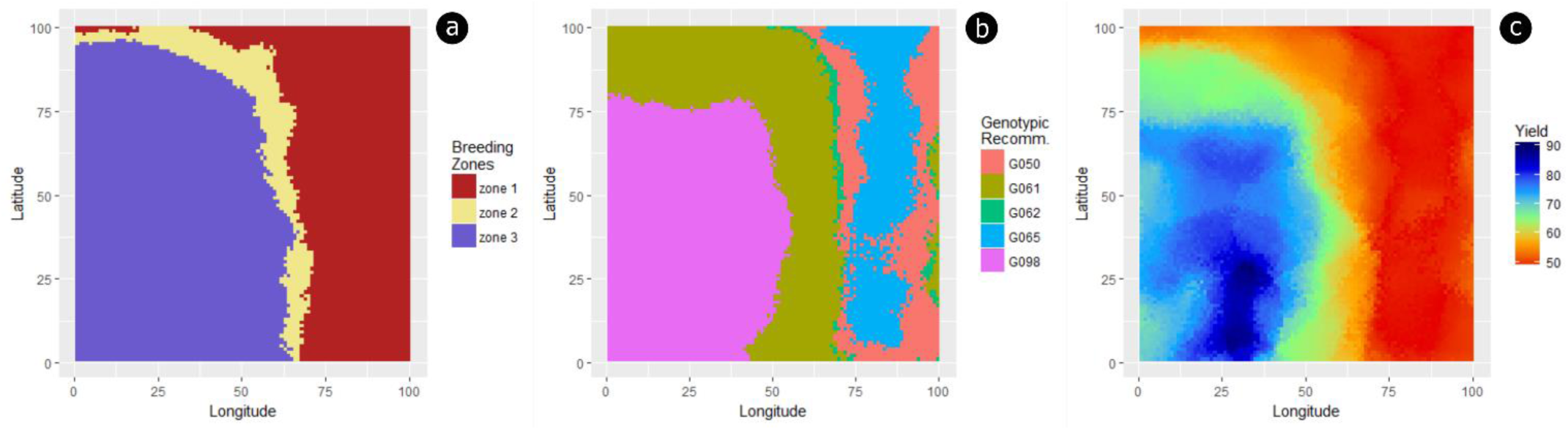
Genetic extrapolations. **(a)** Breeding zones map depicting the three zones fitted by the UPGMA procedure. **(b)** Recommendation map depicting the distribution of the top five genotypes for yield. **(c)** Yield extrapolation for the top five genotypes across the land area. There is a higher genetic correlation (r_g_) within zones, and lower among them, indicating zones with fewer genotypic rank changes.

From the 100 genotypes evaluated through GIS-GE, there were five ranking first in at least one pixel, i.e. G050, G061, G062, G065 and G098, which were best in 14.13, 30.19, 1.85, 15.26 and 38.57% of the whole area, respectively (Figure 4b). When looking for the best genotype for each pixel, the tendency was that neighboring pixels shared the same genotype. When deploying these five recommended genotypes in the field conditions of the evaluated envirome, the expected yield potential is shown in Figure 4c with an average yield equal to 62.24. It is important to note that each pixel in the area has a particular genotypic ranking. For example, when removing the best one (first selected), the second-best genotypes were G006, G032, G041, G074 and G089 in 4.25, 44.93, 25.69, 0.13% and 25.00% of the area, respectively. Thus, different genotypic recommendation panels can be allocated to the area according to the different rankings of genotypes in the environmental gradient.

Considering the two different conditions of unbalanced data, the reduction in the number of experimental trials resulted in a constrained EI predictive ability in recommending genotypes to locations in the area. In terms of yield potential of the genotypes, a reduction in the yield potential of the selected genotypes was observed, but the study indicates that an unbalancing of up to 40% in the total number of trials (a reduction from 50 to ∼20 trials) did not result in great loss of predictive ability for genotype recommendation. However, the evaluation of a very small number of experiments should be avoided, because this condition can, by chance, result in biased yield potential of the area given the use of inadequate genotypes (Figure 5a). In contrast, unbalance of genotypes within the experiments did not present major risks for the application GIS-GE. It should be good practice, however, that a given genotype be allocated in at least 20 of the 50 evaluated experiments (Figure 5b), to ensure an appropriate level of representativeness of the genotypes across the land area.

**Figure 5.**
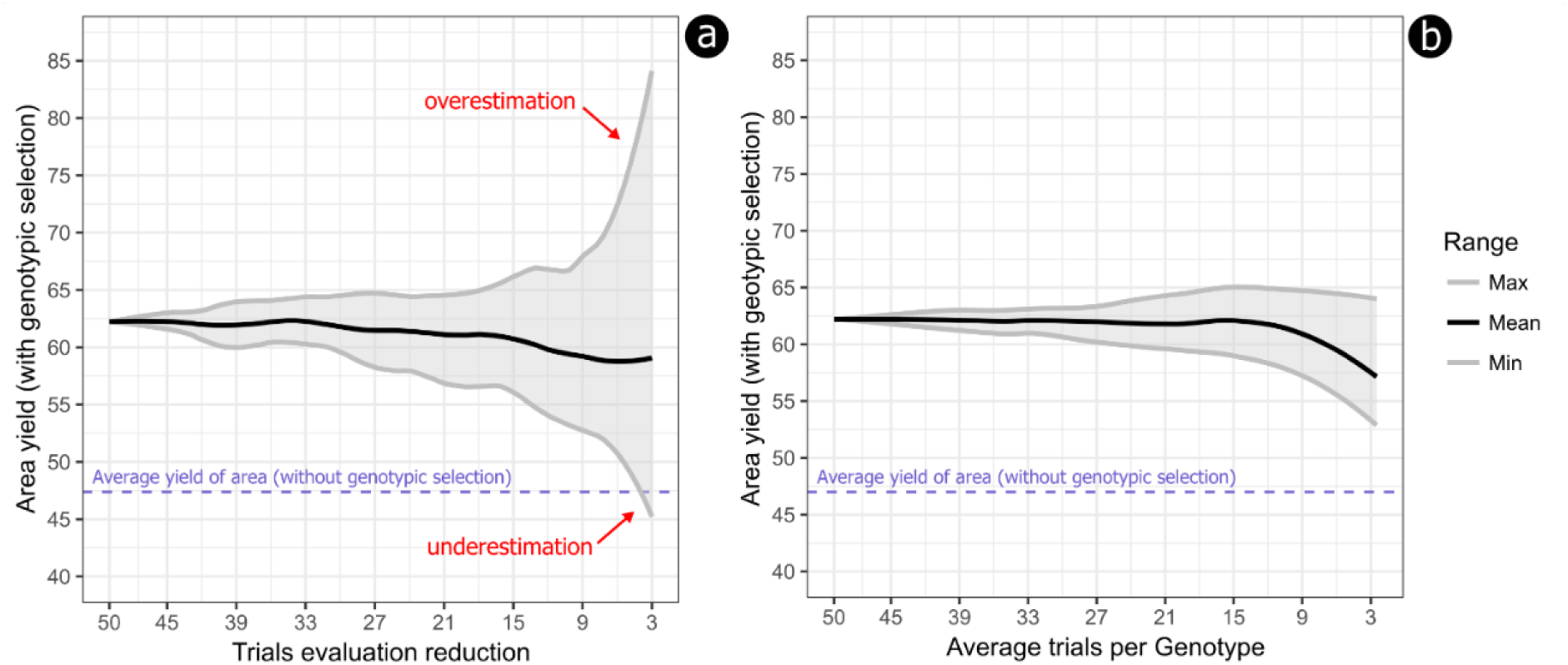
Yield prediction modeling with unbalanced data. **(a)** Impact of reducing the number of trials on yield projection of the selected genotypes. **(b)** Impact of reducing the average number of trials that include a genotype on yield projection of the selected genotypes. The dotted blue line is the average yield for the entire area without genotypic selection/recommendation.

## Discussion

### Enviromics provides a quantitative framework for breeding

In the perspective of maximizing genetic gains through selective breeding, phenotypic traits are thought to be under the influence of several environmental variables. An environmental variable that is favorable for the field deployment of a particular plant species is not always the same for another species. For example, while the presence of exchangeable aluminum in the soil is unfavorable to most annual crops, for some forest tree crops it will be irrelevant (Poschenrieder et al. 2008). In addition, such environmental conditions in a land area may interact at different levels to determine the final genotype yield (Resende et al. 2018). For this reason, the application of the additive main-effects and multiplicative interaction (AMMI) and genotype-and-genotype–environment interaction (GGE) models have been shown, in general, appropriate to detect individual site effects and their interactions on the expression of complex traits (Yan et al. 2000; Gauch 2006). AMMI and GGE regress performance of G×E on multiple latent variables using reaction norms, being natural extensions of the classical regression on the environmental mean, first proposed by Yates and Cochran (1938) and revisited later by Finlay and Wilkinson (1963) and Eberhart and Russell (1966). The regression on the mean is essentially a univariate latent regression, whereas AMMI and GGE are multivariate regressions. If these latter models are reduced to a single multiplicative term, then we are essentially back at the Finlay-Wilkinson regression. But ample experience shows that more than one multiplicative term is needed to properly model the G×E. Our approach rests on the strong assumption that a single latent variable suffices to capture potentially complex G×E. Although this is an optimistic assumption, the enviromic models can be easily adjusted for multiple regressions, for example by grouping the environmental markers into sets with any similarity or by adopting specific objectives. An alternative route that has been followed by others authors is to directly regress genotype responses onto multiple environmental markers (Jarquín et al. 2014; Pérez-Rodríguez et al. 2015), meaning that the interpolation idea can directly be applied in this setting as well.

Finding an environmental index that represents agricultural productivity is one way of characterizing the quality of the environment, independent of the in-depth knowledge of the sources of variation of phenotypic expression. Individually, some environmental factor (an envirotype) may not significantly affect large groups that share diverse levels of genetic relationship across locations in a wide geographical area, but, as a stepwise procedure, may have some influence on the genotype’s performance when combined with other factors in the area (Löffler et al. 2005). For a single envirotype, repeated observation of an individual genotype (or its progeny) may show that its performance is spatially patterned across the locations within the area. In our methodology, the geographical area is an image so that each location is a pixel space in a coordinate system. These coordinates are not very meaningful, but in our data simulation they encoded useful notions of distance (longitude and latitude) and magnitude. Each pixel has intensity in some range, so that the envirotypes are now an image consisting of a big collection of pixels (Figure 1a). Noteworthy is that only few pixels contain genetic trials with phenotypic and possibly genotypic information but all pixels should have environmental information.

Considering all the envirotypes as digital images, we grouped together all similar pixels to account for the effects of all the environmental variables on the genotype’s yield collected from the genetic trials using random forest for classification and regression tasks. In this way, we built an appropriate Environmental Index for the entire land area. The EIs values, rescaled to 0–1, 0 resemble markers that correlate with the environmental suitability and reveal differences on the individual genotype’s performance within the physical area, which we termed enviromic markers.

Grouping together images through their similar pixels is somewhat analogous to the concept of linkage disequilibrium (LD) among genetic markers, the non-random association of alleles at two or more genetic marker loci in a population when dealing with genomic predictive models. It is known that many loci may not be directly involved in the expression of the phenotype itself, but their association to the causal loci due to LD increases the accuracy of predictive models (Jannink et al. 2010). The conceptualization of enviromics in breeding is built on this same principle, that no measurable environmental variable should be neglected, even if in principle it is not found to be statistically related to the phenotype under scrutiny. Like genetic markers spread across the genomes, the combination of EIs may represent the explanation of a representative fraction of the phenotypic variation (Des Marais et al. 2013), either for all genotypes or for some of them.

Analogous to the use of DNA markers, enviromic markers will also require quality control measures, such as Call Rate and MAF (Minimum Allele Frequency). The number of missing-values described by the Call Rate parameter can be solved by adopting two strategies. The first would be to increase the size of all pixels in the area, and use the average information available in the neighborhood. Although this will increase the unit-of-handling area, thereby decreasing the accuracy of the recommendation or prediction, it is an alternative. The second, more feasible, would be to impute the values of the lost pixels by means of kriging modeling, in this case both the neighborhood values and the values of other enviromic markers can be used. For example, similar temperature values between close locations are more likely than distant ones, and on a local scale, the temperature correlates well with terrain elevation, or with global latitude. In the case of MAF, this could be now called MEV (Minor Environmental Variation), the caveat being having environmental covariates with low variance in under sampled pixels compared to all pixels in the area. In this case, enviromic markers with low MEV should be discarded. In the present study, markers were simulated without missing data. Follow up studies will be done to define the optimal thresholds for quality control of enviromic markers.

For breeding practice in the enviromics perspective, the number of field trials in a given land area should satisfactorily support accurate recommendation of genotypes using GIS-GE, i.e., the field trials are expected to cover the range of 0–1 as best as possible, matching the entire range of EI. However, as a continuous random variable, it is very common that the EI values display a normal distribution, so it is also important that the experiments have EI values close to the mean so that a larger area presents good degree of confidence for the application of environmental models and non-linearities can be explored.

Several types of environmental variables can be used in enviromics, such as temporal climatic information (Fick and Hijmans 2017) and Vegetation Indices obtained from remote sensing based canopies (Xue and Su 2017). Xu (2016) lists a number of environmental factors affecting plant growth and yield. Categorical indices can also be used, such as the climatic index of Köppen (Kottek et al. 2006), or even soil classes (Hartemink 2015), since each class is properly represented by one or more experimental points. Although enviromics is expected to be more functional for complex traits, given the intimate relationship between the G × E interaction and quantitative traits, it can also be applied to more discretely distributed traits related to resistance or tolerance to biotic and abiotic stressors, provided that the model for EI generation is fed with data from environmental variables that trigger the targeted physiological stresses. For example, if the focus of the study is improving water deficit tolerance, it is extremely important that data on water availability is included in the envirotyping routine. If the focus is resistance to tick-born diseases in animals, it is important to feed the model with data on the occurrence and density of the arachnids responsible for the disease (Giles et al. 2014). Thus, it is possible to adopt the enviromics approach to traits with different levels of heritability, including disease escape in regions with endemic pests for example (Shakoor et al. 2017).

The accuracy of an enviromic model applied to breeding will depend primarily on the size of the available sample unit (pixel). For instance, when using geographic information, the EI pixel size will delimit the refinement level of the genotypic recommendation (Marcatti et al. 2017), and the level of refinement of breeding areas as well. Environmental information with high spatial resolution, especially resolutions sufficiently adequate to contemplate forest stands, crop plantations and livestock farms, should be preferred to improve the accuracy of the models.

Enviromics may be applied in conjunction with the breeding program strategy specific to each crop or animal or at any specific stage of the program. Annicchiarico and Iannucci (2008) suggest genetic improvement pathways specific to each geoclimatic area based on distinct genetic basis and selection environments. In maize breeding, for example, differences in the capture of genetic variances can be observed along the trials carried out throughout the breeding program (Cooper et al. 2014). Enviromics may also have applications in recurrent selection programs, directing preferred crosses and selecting suitable parents. One can take advantage of the predicted pixel-by-pixel genetic values by coupling kinship structures to the enviromics models, according to the stage of the breeding program and simply devise the strategy as best suited. One can also perform it in a mixed model by forming the direct product of the Kinship matrix (*A*) with the reaction norm covariance (Σ_a_). In fact, this could be stated explicitly with the mixed model, it’s *A* ⨂ Σ_a_, where ⨂ is the kronecker product.

Additionally, fundamental to the program is the generation of phenotypic variability, linked to the use of genetic resources from germplasm banks. The enviromics models can also direct the rescue of materials with local-specific phenotypes, or according to the local consumer profile, directing the trials to target populations of environments (TPE) (Chapman et al. 2000). Our TPE, with all pixels of a rectangular area (a naïve strategy), it is only an illustration, which could obviously display a much larger domain, such as the combined use of on-farm data from an entire country, sets of countries or data collected across countries, by which economically underdeveloped countries could benefit (Pérez-Rodríguez et al. 2017). In other words, a TPE can be quite large, depending on data availability. Phenotypic data should only be linked to the crop growth period. After that, we can adjust the models for scenarios of climate change or associate them with complex geospatial weather forecast data. It is important to note that while the breeder does not need to have an in depth understanding of geotechnologies, a reasonable assumption is that the breeder should get acquainted with the use of spatial data.

From the point of view of genetic selection, a great benefit of enviromics models is the possibility of quantifying residual variances inherent to the whole geographic environment, and possibly maximizing the genetic variance captured especially in those steps that display low heritabilities. Thus, in addition to maximizing the genetic variance components for the selection of individuals, families or parents (Gomez-Raya and Burnside 1990), it is possible to optimize selection by identifying sites that provide a higher ratio between genetic and residual variances. Especially in populations with a high level of improvement, or even in the stages of the program when trait heritabilities are lower, such high-heritability environments can help increasing the accuracy of genotypic selection.

### Case Study: Recommendation of genotypes using GIS-GE

In the routine of a crop breeding program, it is common practice to allocate experiments in several sites throughout a land area, to cover better the spectrum of possible sites of genotype recommendation (Annicchiarico et al. 2006). This procedure aims to identify those particular genotypes that may be recommended for the largest number of sites in terms of their phenotypic means in comparison to their competitors. Underlying this practice, the assumption is that the top genotypes in the evaluated trials may perform better across the whole area where the breeding experiments were deployed. The extrapolation arguments are mostly informal and unquantified because they rely on parameters that are assumed true beyond the range of values for which they are known to be true. It can be noted, thus, that these practices often carry implicit arguments in favor of the notion that the evaluated and non-evaluated sites across the land area show a high level of correlation among them in terms of the influence of their environmental variables on growth and yield. The high correlations assumed among every possible variable considered within every possible site across the area are deemed sufficient to preclude changes in genotypic ranking.

In contrast to this canonical view, we have introduced the potential application of enviromics in genetic improvement, by presenting the GIS–GE methodology to select and recommend genotypes for non-sampled sites. For this purpose, a network of genetic trials was simulated in the final stage of the breeding program such that genotypes in these trials could be hybrid varieties or commercial lines, vegetatively propagated clones, or even animal breeds. In our simulation, the environmental variability was considered a preponderant factor in the generation of phenotypic variability. Therefore, one hundred different environmental variables were adopted arbitrarily, without necessarily correlation existing between them (Figure 1a). As a consequence, the results indicate that ranking of genotypes may vary abruptly among EI across the land area, even for those which display very similar environmental variability depicted in terms of their corresponding EI (Figure 2b). The genotype with the greater phenotypic mean (G061), which was recommended for approximately 30% of the area was not even close, in terms of yield, to genotype G050 ranked first in most trials (Figures 2b and 4b). In addition, the analysis indicated a poor performance of G061 in environments with EI <0.25 (Figure 2b). An inverse interpretation can be made for genotype G039, which was the worst in terms of phenotypic mean (Figure 2b). Another interesting case was genotype G074, which showed low phenotypic values in environments with EI> 0.25. Although it may be a bad recommendation for environments with higher EI, it demonstrated good performance in environments with low productive potential, and could be recommended for these locations in some situations.

Models based on reaction norms are interesting because they show the stability of genotypes to environmental changes and their adaptive ability. Although stable genotypes may be attractive, as they generally do not result in unexpected surprises to the breeder, adaptive genotypes may respond better to crop management procedure such as irrigation and fertilization (Cobb et al. 2013), control of biotic agents (Shakoor et al. 2017) or even improvements in animal comfort (Reynolds 1996). When observed, negative predicted yields in lower EI may indicate that the genotype would not survive in such environments, or would have exceedingly low yields. On the other hand, genotypes with positive predicted yields in extreme environments may indicate genotypes with high resilience potential. These resilient genotypes usually do not have high regression slopes and are considered to be less adaptive and very stable on their response to different environmental variables (Chloupek et al. 2004) (Figure 2c).

Although simulations were carried out with a low heritability trait, results showed that it is possible to capture greater heritabilities in extreme locations (EI≥0 or EI≤1) (Figure 3a). Although Bänziger (2000) argues that differences between genotypes are generally smaller under stress and larger differences are, therefore, more difficult to detect, sites with limiting characteristics to plant development may lead to selection pressure on the most adapted individuals (McKown et al. 2014), favoring the manifestation of more productive genotypes. Sites with better conditions may also provide similar results, since some genotypes may demonstrate better ability to take advantage of the available resources than others. This behavior is usually demonstrated in reaction norm studies in animal breeding (Santana Jr et al. 2013; Ambrosini et al. 2014), forest tree improvement (Resende et al. 2018), and agricultural grain crops (Jarquín et al. 2014).

Identifying the spatial boundaries in which a selected genotype may be recommended would be an important advance (Annicchiarico et al. 2005), considering that it is usually impossible to deploy a large number of trials in a certain area due to limited resources. Across the EI built with the GIS-GE method, our results showed genetic correlations between 1.00 and -0.49, indicating that depending on the relationship between selection and recommendation site, it is possible to hit with absolute certainty, or to make a major mistake (Figure 3b). The efficacy of a system of evaluation of cultivars depends largely on the genetic correlation between genotype performance in multi-environmental trials (MET) (Löffler et al. 2005). A map of breeding zones is therefore indicated, precisely linked to the size of the pixel used (Figure 4a). Three breeding zones were observed in our simulations, with high genetic correlations within each one, indicating high confidence between the selection and recommendation site. When no trials are available in one particular breeding zone, one can use selections of a zone with the highest average correlation. In our work the genetic correlation between the khaki and blue zones was 0.70, indicating that no drastic G × E interaction would be expected between these two macro-environments. Furthermore, within a breeding area, priority should be given to allocate trials where a better ability to capture missing heritability is expected, as sites with higher heritability are prone to be sites of higher selective accuracy (Marcatti et al. 2017).

With a recommendation map in hands, the breeder or the agricultural extension service may indicate better genotypes for very specific boundaries in the area (Figure 4b). More than one genotype should be recommended per site, contemplating two points highlighted by Annicchiarico et al., (2006): (*i*) mitigate the risk of unexpected susceptibility to a biotic or abiotic stress by a single recommended cultivar; (*ii*) take into account genotypic differences that were not statistically significant during recommendation analysis. Schedule and logistic issues can also be incorporated into enviromics models, such as the availability of improved seeds for grain crops or seedlings for forest trees. If the recommended genotypes are effectively deployed in the area, the potential yield may increase by 32% when compared to the average yield of the area planted with unselected genotypes (Figure 5). This is a key point as it meets the challenge of increasing production with the same land area.

The proposed genetic modeling adopted in the GIS-GE methodology is based on mixed models, which deliver a powerful statistical framework for dealing with unbalanced data (Gilmour et al. 1995). However, when tested with unbalanced dataset, by radically reducing the number of experiments in the field, the ability of the models to select genotypes that maintained the average yield was not satisfactory. Although mixed models have the ability to work with unbalanced data, like any other statistical procedure, they are unable to correct for exceedingly inefficient sampling (Schmidt et al. 2019). On the other hand, genotypic imbalance was not detrimental to the recommendation ability of the GIS-GE model. In practice, these results indicate that if a choice has to be made, it is better to establish more and smaller trials with less replication of genotypes than deploying a smaller number of replicated trials (Moehring et al. 2014). An insufficient number of trials may sometimes over- or underestimate the yield of selected genotypes. This happens because when the trials show yields above the average of the area, the yield of the recommended genotypes will be overestimated in relation to the expected future yield. The reverse is also true when an inadequate sampling of trial environments happens that result in below-average yields. While Pérez-Rodríguez et al. (2015), evaluating only nine experiments, achieved predictive capacity gains of approximately 4-6%, Jarquín et al. (2014) achieved up to 34% gain by evaluating on-farmer trials addressing 134 locations.

### Future implications of enviromics in breeding

The convergence of genomics and quantitative genetics is currently revolutionizing plant improvement especially for long lived perennial plants where genomic prediction is particularly appealing to accelerate breeding cycles (Grattapaglia et al. 2018). Our work provides a glimpse into the promising area of further including enviromics in this context to optimize the ability to predict the performance of breeding material, especially for species subject to complex response patterns across environments and time. The enviromics models presented may seamlessly incorporate any genetic structure by changing the existing relationship matrix between genotypes from the traditional numerator ***A*,** to the genomic ***G***, or combined A and G based on H matrices. This last one is a hybrid matrix using both kinship information and molecular marker data, was shown to increase by 3-5% the accuracy of breeding models (Legarra et al. 2009). In the context of using ***G*** or ***H*** matrices, improved genomic predictions of complex phenotypes are expected across environments by simultaneously taking into account information from molecular (SNP marker data) and enviromic markers, especially for late-expressing or difficult to measure phenotypes. Additionally, given the very high genotyping density possible with current SNP panels, the enviromics approach may also be used to generate a catalog of SNP markers for each micro-region in a target area, naturally respecting the strong G × E typically displayed by traits of low heritability. Finally, genomic-based enviromics models can also exploit precise field-level information of the trial, such as competition between plants (Cappa et al. 2017). Additionally, environmental information can be incorporated the prediction models via envirotyping (Xu 2016) combining genomics, controlled crosses, germplasm data and next-generation phenotyping (Cobb et al. 2013). We can also mention that epigenetic effects expressed triggered by changes in the environment can also be better captured with enviromic models, by the incorporation of epigenetic matrices ***T*** into the breeding analysis (Varona et al. 2015).

The construction of environmental indexes can also benefit from the application of artificial intelligence, eco-physiological process models (Asseng et al. 2013) or biogeographical similarity approaches (Vilhena and Antonelli 2015), in order to achieve indexes that relate more closely to the target trait, avoiding the inclusion of trait-irrelevant enviromic markers. As the methodology maximizes the number of recommended genotypes based on the potential yield of the area, one should also be concerned with the maintenance of genetic variance throughout the selection cycles. This can be done using optimization models that maximize selection gains with genetic diversity of the selected individuals (El-Kassaby and Lstibůrek 2009; Mullin 2017). Another interesting strategy would be the recommendation of specific parents and crosses for specific environments, generating progeny based on the combination of the best individuals by environment (van Ginkel and Ortiz 2017).

A possible limitation of the proposed GIS-GE method is the assumption of a common G × E profile for two sites with similar environment index. Different factors that make the two sites similar for productive potential may in fact result in different G × E patterns. To overcome this situation, a single-step enviromics model would improve the prediction accuracy over environments. These models can be purely Bayesian, since covariates associated are treated as unknown, thereby allowing inference for all unknowns together within a one-step linear random regression as follows: *y* = ***X****β* + ***P****w* + ***Z***_**0**_*a*_0_ + ***Z***_**1**_*a*_1_ + *e*, where: *y* is the vector of observations, *β* is the fixed effects vector of order *p*, 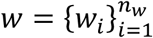 is the vector of environmental effects, 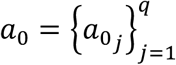 is the vector of random genetic intercepts, 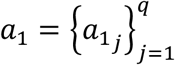 is the vector of random genetic slopes. It can be assumed that: [*a*_0_, *a*_1_]^′^∼N(0, ***G*** ⊗ ***Σ***_*a*_), being ***G*** the genomic relationship matrix under a GBLUP framework. Furthermore, ***X***, ***P***, ***Z***_**0**_ and ***Z***_**1**_ are the respective known incidence matrices, whereas the row address of ***Z***_**1**_ has exactly one element equal to the effect of the environmental covariate (*w*_*i*_ or an estimate of *w*_*i*_), with all other elements in that row equal to zero, and *e* is the vector of random residuals. To infer environmental sensitivities, three stages are required: the first stage defines the distribution of the phenotypic data conditional on all other parameters; the second stage is represented by the prior distributions of the location parameters *(β, w, a*_0_, and *a*_1_); and the third one is based on specifying prior distributions for the (co)variance components.

In addition, the previously described model is based on GBLUP, which is not suitable for variable selection (shrinkage estimates) in the presence of many correlated covariates (enviromic or molecular markers). However, we think that other specific genome prediction models like Bayes A, B or Bayesian LASSO can be adapted to infer on G × E. For example, for these three Bayesian regression models, the following prior distributions for the effect of marker i at environment level j (α_ij_) could be adopted assuming heterogeneity of marker effect variances over all environment levels: 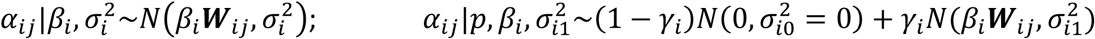; and 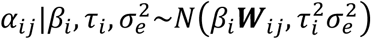, where in all distributions *β*_i_ is the slope of the SNP effect associated to environment level ***W***_*ij*_. It is clear that under these approaches, the Bayesian modeling needs more than one step in its hierarchy, which is given by the assumption of an appropriate prior distribution for *β*_i_.

## Concluding remarks

The term *enviromics* in the context of breeding practice involves the application of envirotyping techniques to describe the performance of a plant or animal along the different gradients of a large number of environmental variables. To account for the effects of environmental variables on a phenotype we have developed an infinitesimal-like model taking into account additive and non-additive contributions from enviromic markers in an analogous fashion as a quantitative genetics model. Multivariate models, such as principal components analysis, and modern approaches from artificial intelligence will likely allow better definition of enviromic markers improving the computation of Environmental Indexes. Enviromics models are flexible and can be easily adjusted according to changes in the environment, a particularly useful feature in the context of climate change scenarios. Additionally, enviromic markers are climatic or landscape-based variables. As such, not only are they universal for any animal or plant species but more importantly they can be obtained in a significantly cheaper and faster way than any other omics marker data such as DNA, RNA, proteomic or epigenomics. Finally, we have also proposed a methodology called GIS-GE derived from the enviromics conceptualization, to recommend genotypes for specific areas, to define optimal breeding zones, to understand the spatial boundaries in which a genetic trial can be used for selecting breeding material, and to identify sites that provide better capabilities for the genetic expression of a phenotypic trait. We trust that the concept and methodology presented here should represent a relevant advance in the existing approaches and be a useful addition to the toolbox of the modern breeder, especially with the increasing availability and use of genomic and environmental big data.

## Author contribution statement

RTR, HPP, OBSJ and FFS: simulation and data analysis. MDVR, RTR, HPP and FFS developed mathematical-statistical procedures and notations. RTR, HPP, OBSJ, FFS, MDVR and DG: wrote the manuscript. All authors reviewed the manuscript.

## Acknowledgments

Professors Gustavo E. Marcatti from UFSJ and Helio G. Leite from UFV for important plant management and GIS concerns; *Fundação de apoio à pesquisa do Distrito Federal* (FAPDF); and the National Council for Scientific and Technological Development (CNPq) for funding resources.

## Availability of data and materials

Code used to generate the simulated data, and the envirotypic (File S1) and phenotypic data (File S2) are available from us in the repository: https://figshare.com/articles/Enviromics_data/8264132.

